# A study of the correlation between phenotypic antimicrobial susceptibility testing methods and the associated genotypes determined by whole genome sequencing for a collection of *Escherichia coli* of bovine origin

**DOI:** 10.1101/2021.07.16.452668

**Authors:** Thomas J Maunsell, Scott Nguyen, Farid El Garach, Christine Miossec, Emmanuel Cuinet, Frédérique Woehrlé, Séamus Fanning, Dagmara A. Niedziela

**Affiliations:** UCD-Centre for Food Safety, UCD School of Public Health, Physiotherapy and Sports Science, University College Dublin, Belfield, Dublin D04 N2E5, Ireland; Department of Biopharmaceutical and Medical Science, Galway-Mayo Institute of Technology, Dublin Road, Galway, Ireland; Microbiology Laboratory, Pathology Department, Blackrock Clinic, Rock Road, Blackrock, Dublin, Ireland; Public Health Laboratory Division, District of Columbia Department of Forensic Sciences, Washington, District of Columbia, 20024, USA; Vétoquinol SA, BP 189-7024 Lure, France; School of Veterinary Medicine, University College Dublin, Belfield, Dublin 4, Ireland

**Keywords:** Antimicrobial resistance, *Escherichia coli*, disk diffusion, minimum inhibitory concentration, bovine isolates, genotype-phenotype correlation, whole genome sequencing

## Abstract

Antimicrobial resistance (AMR) has increased at an alarming pace in the recent years. Molecular-based methods such as whole genome sequencing (WGS) offer a potential alternative to the conventional labour-intensive methods traditionally used to characterise AMR phenotypes. The aim of this study was to investigate whether WGS could be used as a predictor of AMR in *Escherichia coli* isolates of bovine origin.

Genomes of 143 *E. coli* cultured from cattle presenting with diarrhoea or mastitis were sequenced on an Illumina MiSeq platform. AMR genes were identified using the ResFinder and AMRFinder databases. Antimicrobial susceptibility testing by disk diffusion was performed on a panel of 10 antibiotics, covering 7 antimicrobial classes. Minimum inhibitory concentration (MIC) measurements were made using the Sensititre plate with 6 antibiotics, covering 5 antimicrobial classes. Correlation between genotype and phenotype was assessed statistically by means of a two-by-two table analysis and Cohen’s kappa (κ) test.

The overall κ correlation between WGS and disk diffusion was 0.81, indicating a near perfect agreement, and the average positive predicted value was 77.4 %. Correlation for individual antimicrobial compounds varied, with five yielding near perfect agreement (κ = 0.81–1.00; amoxicillin, florfenicol, gentamicin, tetracycline and trimethoprim-sulfamethoxazole), one showing substantial agreement (κ = 0.65; nalidixic acid), and four showing moderate agreement (κ = 0.41– 0.60). The overall κ correlation between WGS and MIC was 0.55 indicating moderate agreement, and the average positive predicted value was 68.6 %. Three antibiotics yielded near perfect agreement (gentamicin, tetracycline and trimethoprim-sulfamethoxazole) and a further three showed fair agreement (κ = 0.21–0.40).

WGS is a useful tool that can be used for the prediction of AMR phenotypes, and correlates well with disk diffusion results. MIC measurements may be necessary for antimicrobial compounds with a high proportion of intermediately resistant isolates recorded, such as cephalothin.

**Highlights:**

- Culture based antimicrobial susceptibility testing is used to identify therapeutics in the treatment of clinical veterinary isolates
- Whole genome sequencing is increasingly adopted for surveillance, epidemiological traceback investigations, and detection of antimicrobial resistance genes
- Little is known in correlations between antimicrobial resistance genotypes and disk diffusion antimicrobial susceptibility testing
- This study finds that whole genome sequencing is a useful predictor for antimicrobial susceptibility however, minimum inhibitory concentration measurements may still be needed for intermediately resistant isolates

## Introduction

Antimicrobial resistance (AMR) is a modern societal challenge with veterinary public health dimensions. This situation has arisen owing to the widespread use of antimicrobial compounds in various settings to treat human and animal infections (Zankari et al., 2013). Emergence of AMR in bacteria of importance to veterinary public health has consequently limited the chemotherapeutic options available for effective treatment of clinical infections. In the case of some multi-drug resistant (MDR) bacteria this development has given rise to the fact that no treatments are available, for instance for carbapenem-resistant *Enterobacteriaceae* (CRE) which are associated with human nosocomial infections globally (Jasovský et al., 2016). In 2015, over 670,000 antibiotic resistant infections were recorded in Europe with approximately 33,000 associated deaths thereby highlighting the need for rapid and accurate antimicrobial susceptibility testing which offers clinical utility (Cassini et al., 2019). AMR can evolve through several mechanisms including point mutations in target housekeeping genes, or *via* horizontal gene transfer (HGT) of AMR-encoding genes mapped to various mobile genetic elements (MGEs) such as plasmids, transposons and prophages. Upregulation of gene expression for efflux pumps can also contribute to this resistance phenotype in bacteria (Punina et al., 2015). The continual evolution of multidrug resistance highlights the fact that AMR will continue to pose a challenge for veterinary public health in the future (Feldgarden et al., 2019).

Antimicrobial susceptibility testing (AST) is an important laboratory method for determining which antimicrobial compounds will exhibit therapeutic success if administered to treat a bacterial infection. To reduce the inappropriate use of these drugs whilst attempting to limit the subsequent development and spread of antibiotic resistant bacteria, AST should be performed to guide clinical treatment decisions. Susceptibility testing is essential for detecting the emergence of new resistance phenotypes, determining the extent of AMR in bacterial populations and assessing changes in resistance patterns over time (Zankari et al., 2013). Traditional AST relies on phenotypic-based methods that are cost effective and sparing in terms of resources. Molecular-based methods are also available and based on the amplification of a known gene or more recently the application of whole genome sequencing (WGS). Using WGS, it is possible to detect all AMR-associated genotypes in a given bacterium, rather than evaluating a selected panel of antimicrobial agents with traditional laboratory-based methods. In order to understand AMR fully and improve AMR diagnostics, its link to bacterial genotype must be further understood (Feldgarden et al., 2019). WGS also has the potential to extrapolate new information, e.g., the presence of novel antimicrobial resistance genes, from bacterial genomes, which may alter how future antimicrobial resistance testing is performed.

WGS is currently a valuable diagnostic tool applied to outbreak investigation, tracking and surveillance (Ribot et al., 2019). The potential of WGS to aid in the detection of AMR is substantial, allowing for detection of novel resistance variants (Feldgarden et al., 2019). Increased accessibility to genomic databases *via* electronic means along with a reduction in the cost of sequencing can, facilitate the replacement of those traditional methods, in time with WGS in the clinical veterinary laboratory (Tyson et al., 2015; Moran et al., 2017). WGS-based AMR genotype detection is a rapidly developing area with the potential to accurately and rapidly predict all known resistance genotypes for an individual isolate using a single assay (Su et al., 2019). This gives WGS-based methods a clear advantage over culture-based AST methods and PCR where the number of resistant phenotypes determined is dependent on the antimicrobial resistance-encoding genes selected for inclusion (Su et al., 2019). Once generated, WGS data can be used for other purposes, such as investigation of virulence or phylogeny determination. This approach aims to predict the antimicrobial resistance phenotype of a bacterium that would otherwise have been obtained if the isolate was tested phenotypically for the same trait. This is done by cross referencing the sequence obtained following WGS, using a suitable bioinformatics pipeline, with a database of known resistance genes such as ResFinder, CARD or AMRFinder (Zankari et al., 2012; McArthur et al., 2013; Feldgarden et al., 2019). AMR genotypes can be predicted based on the presence of these AMR genes or mutations in highly conserved peptides which are known to play a key role in bacterial cell physiology (Su et al., 2019). Mutations also occur in accessory genes, which code for mechanisms such as efflux pumps, which results in antibiotic resistance (Vogwill and Maclean, 2015). Studies that compare phenotypic and genotypic resistance profiles are needed in order to establish the reliability of WGS-based approaches as a stand-alone predictor of true AMR in bacteria (Moran et al., 2017).

A number of studies have been performed to assess the correlation between WGS and phenotypic AMR (Card et al., 2013; Stoesser et al., 2013; Zankari et al., 2013; Tyson et al., 2015; Feldgarden et al., 2019; Stubberfield et al., 2019). Reported correlation study data, however, can depend on the nature of the antibiotic; the bacterial species tested and the associated mechanism of resistance. The cited studies use MIC as the phenotypic AST method of choice for comparison. Little is known about the correlation between disk diffusion AST and WGS antimicrobial resistance testing. Therefore, further research evidence is required to determine whether WGS can accurately infer antimicrobial susceptibility (Stubberfield et al., 2019).

*Escherichia coli* is found as part of normal gut flora of both humans and warm-blooded animals. Most *E. coli* are non-pathogenic but can act as a reservoir of AMR genes, which can be transmitted to other bacteria sharing the same ecological niche thereby posing a threat to human health (Stubberfield et al., 2019). Transmission to humans occurs through consumption of undercooked or contaminated animal products such as hamburgers or milk. Transmission can also occur through consumption of plant-based foods contaminated with faecal matter, and direct contact between livestock and farmers (Espinosa et al., 2018). This emphasises the need for a multisectoral One Health approach to combating antimicrobial resistance.

The aim of this study is to investigate the use of WGS as a modern diagnostic tool to accurately predict AMR phenotypes in *E. coli* of bovine origin. Genotypic and phenotypic AMR results of 143 *E. coli* of bovine origin were compared to determine the correlation between the resistance genotype and the phenotypic expression of the determinant.

## Methodology

### Bacterial Isolates and culture methods

A total of 143 *E. coli* of bovine origin were isolated from faecal and milk samples taken from animals presenting with mastitis and diarrhoea. These cattle were located in Germany and France and samples were obtained between the years 2004 and 2010 **(Table 1).**

**Table 1:** A table showing the breakdown of test isolates (n=143) by country of origin, disease and year isolated

| Country of origin |  | Disease |  | Year of isolation |  |
| --- | --- | --- | --- | --- | --- |
| France | 67 | Mastitis | 70 | 2004 | 48 |
| Germany | 76 | Diarrhoea | 73 | 2007 | 48 |
|  |  |  |  | 2010 | 47 |

Isolates were placed in Lysogeny Broth (LB) (Sigma Aldrich) with 15 % [v/v] glycerol and stored at −80 °C. Working cultures of each isolate were prepared by streaking the LB broth onto Mueller-Hinton (MH) agar (Sigma Aldrich) and from there taking three single colonies and re-streaking onto MH agar. Three single colonies from the second inoculum were used to inoculate 5 ml MH broth (Sigma Aldrich). The MH broth was incubated overnight at 37 °C, with orbital shaking at 200 rpm to promote growth.

### Whole Genome Sequencing

#### DNA isolation, purification and library preparation

Total genomic DNA (gDNA) from overnight cultures of bacterial isolates grown in MH broth was purified using the DNeasy UltraClean Microbial Kit (Qiagen). Quality and quantification of total gDNA extract was assessed using the NanoDrop Microvolume Spectrophotometer (Thermo Fisher Scientific). Following quantification, total gDNA was serially diluted 1:10 for each isolate to a concentration of 2 ng/μl which was confirmed by fluorometric quantification using the Qubit fluorometer and a dsDNA High Sensitivity assay kit (Thermo Fisher Scientific). Libraries were generated using the Nextera XT Library Preparation Kit (Illumina). All libraries were then subjected to 300 bp pair-end sequencing (V3 chemistry) on a MiSeq sequencer (Illumina) to produce raw read data.

#### Whole genome assembly and annotation

FastQC (v0.11.7) (Wingett and Andrews, 2018) was used to assess raw read quality while Trimmomatic (v0.39) (Bolger et al., 2014) was used to trim low-quality sequences. SPAdes (v3.13.0) (Bankevich et al., 2012) was then used to *de novo* assembly trimmed paired reads and the quality of the resulting contigs was determined using QUAST (v5.0.2) (Gurevich et al., 2013).

#### Identification of AMR genes, serogroup, phylogroup and strain type

Identification of AMR encoding genes was carried out using Abricate (v1.0.0; https://github.com/tseemann/abricate) and the ResFinder databases (version last updated on 14 May 2020; (Zankari et al., 2012), as well as using NCBI AMRFinder Plus (v3.8.4; Feldgarden et al., 2019). The criteria for defining AMR gene presence included a cut-off of 80 % for gene coverage and 95 % for gene identity. A consensus from both ResFinder and AMRFinder databases was used for final genotype to phenotype correlation. Multi-locus sequence typing (MLST) was performed on the isolates using the mlst package (v2.18.1; https://github.com/tseemann/mlst). Serotyping was carried out using the EcOH database within Abricate.

#### Phylogenomic analysis

Enterobase (https://enterobase.warwick.ac.uk/) was used to curate strains from the whole genome sequencing project (BioProject Accession PRJEB32666) (Zhou et al., 2020). A SNP project in Enterobase (Zhou et. al, 2020) was used to construct a phylogenetic tree. In brief, *E. coli* genomes were aligned against reference CFS3246 using default parameters in the Enterobase SNP project. 313947 variant sites were called with a minimum of 95% sites present in all genomes. The tree was visualised in GrapeTree in Enterobase (Zhou et. al., 2020). Phylogroup prediction was also performed within Enterobase. Additional metadata and sample IDs are available in Table S1.

### Data availability

Fastqc files are available in European Nucleotide Archive (ENA), project number PRJEB32666. https://www.ebi.ac.uk/ena/data/view/PRJEB32666

Genome assemblies and associated metadata are available in Enterobase under Bio Project ID: PRJEB32666. http://enterobase.warwick.ac.uk/species/ecoli/search_strains

### Antimicrobial Susceptibility Testing by Disk Diffusion (AST-DD)

Antimicrobial Susceptibility testing was carried out on all 143 isolates in accordance with Clinical and Laboratory Standards Institute (CLSI) guidelines for disk diffusion. A total of 10 antibiotics were used covering 7 antimicrobial classes and included: amoxicillin (10 μg), amoxicillin/clavulanic acid (20/10 μg), gentamicin (10 μg), cephalothin (30μg), ciprofloxacin (5μg), florfenicol (30μg), marbofloxacin (5μg), nalidixic acid (30μg), tetracycline (30μg), trimethoprim-sulfamethoxazole (1.25/3.75μg). Each antimicrobial was tested in triplicate and the average zone size was used to determine resistance/sensitivity according to CLSI document VET01S, 5^th^ ed. *E. coli* ATCC™25922 was used as a control strain.

### Antimicrobial Susceptibility Testing by minimum inhibitory concentration (AST-MIC)

Minimal inhibitory concentrations (MIC) of 22 antibiotics against 82 *E. coli* isolates was determined using the ready to use Sensititre Gram Negative MIC Plates GN3F (ThermoFisher Scientific) according to manufacturer’s instructions. Briefly, a 0.5 McFarland suspension in physiological water was prepared from an agar overnight culture of each isolate. Then the suspension was diluted in cation adjusted Mueller Hinton broth to reach 10^5^ CFU/mL. 50 μL of diluted inoculum was then used to inoculate each well of the Sensititre plate except negative control wells and the plate was incubated at 34-36°C for 18-24 hours. MIC results were interpreted according to CLSI document VET01S. Antibiotics were selected for correlation if 10 or more phenotypically resistant isolates and AMR genes were present for a given compound, and included ampicillin, cephalothin, ciprofloxacin, gentamicin, tetracycline and trimethoprim-sulfamethoxazole.

### Statistical analysis

Correlations between genotypic and phenotypic data obtained in this study was assessed statistically by means of a two-by-two table analysis (Mackinnon, 2000). AMR gene presence/absence (obtained from WGS data) was compared with phenotypic disk diffusion/MIC AST data giving the actual resistance profile of the isolate. Sensitivity, specificity, positive predicted value (PPV) and negative predicted value (NPV) were calculated taking the following into consideration:

- True positive – gene present and disk diffusion/MIC resistant
- False positive – gene present and disk diffusion/MIC susceptible
- True negative – gene absent and disk diffusion/MIC susceptible
- False negative – gene absent and disk diffusion/MIC resistant

Based on recommendations for surveillance of antimicrobial resistance, intermediate isolates were treated as resistant and subsequently referred to as “intermediately resistant” (Cornaglia et al., 2004). Cohen’s kappa (κ) test was performed to measure the agreement of AST-WGS to Disk Diffusion/AST-MIC in the detection of AMR. A κ value of 1 would indicate perfect agreement.

## Results and Discussion

The bovine *E. coli* cultured from diarrhoea and mastitis cases and used in this study were sequenced using short-read sequencing on an Illumina MiSeq platform and antimicrobial resistance genes were identified by bioinformatic analysis. Phenotypic AST by disk diffusion (AST-DD), including 10 antimicrobial agents, covering 8 classes, was carried out and standardised to CLSI guidelines (CLSI, 2021). A subset of 82 isolates with more than one AMR gene was subsequently selected for MIC analysis using Sensititre plates (AST-MIC). A total of 6 antibiotics belonging to 5 antimicrobial classes were used for correlation between genotype and MIC.

### Population structure of French and German bovine-associated *E. coli*

During 2004-2010, 147 *E. coli* were isolated from animals presenting with diarrhoea or mastitis symptoms and subjected to whole genome sequencing, with 4 genomes excluded from the analysis (CFS3241, CFS3306, CFS3348, and CFS3367) due to failed genome assembly QC. The genomic features of these isolates have been described elsewhere (manuscript in submission, 2021).

The isolates belonged to phylogroups A (n = 45), B1 (n = 42), B2 (n= 5), C (n = 18), D (n = 20), E (n = 11) and G (n = 2), as determined by ClermontTyping in Enterobase (Clermont et al., 2000, 2013; Beghain et al., 2018) (**Fig. 1A**). Bovine mastitis isolates were broadly distributed throughout all phylogroups, supporting evidence that there are no clear related genotypes for a proposed bovine mastitis pathotype (Leimbach et al., 2017) (**Fig. 1B**).

**Figure 1:**
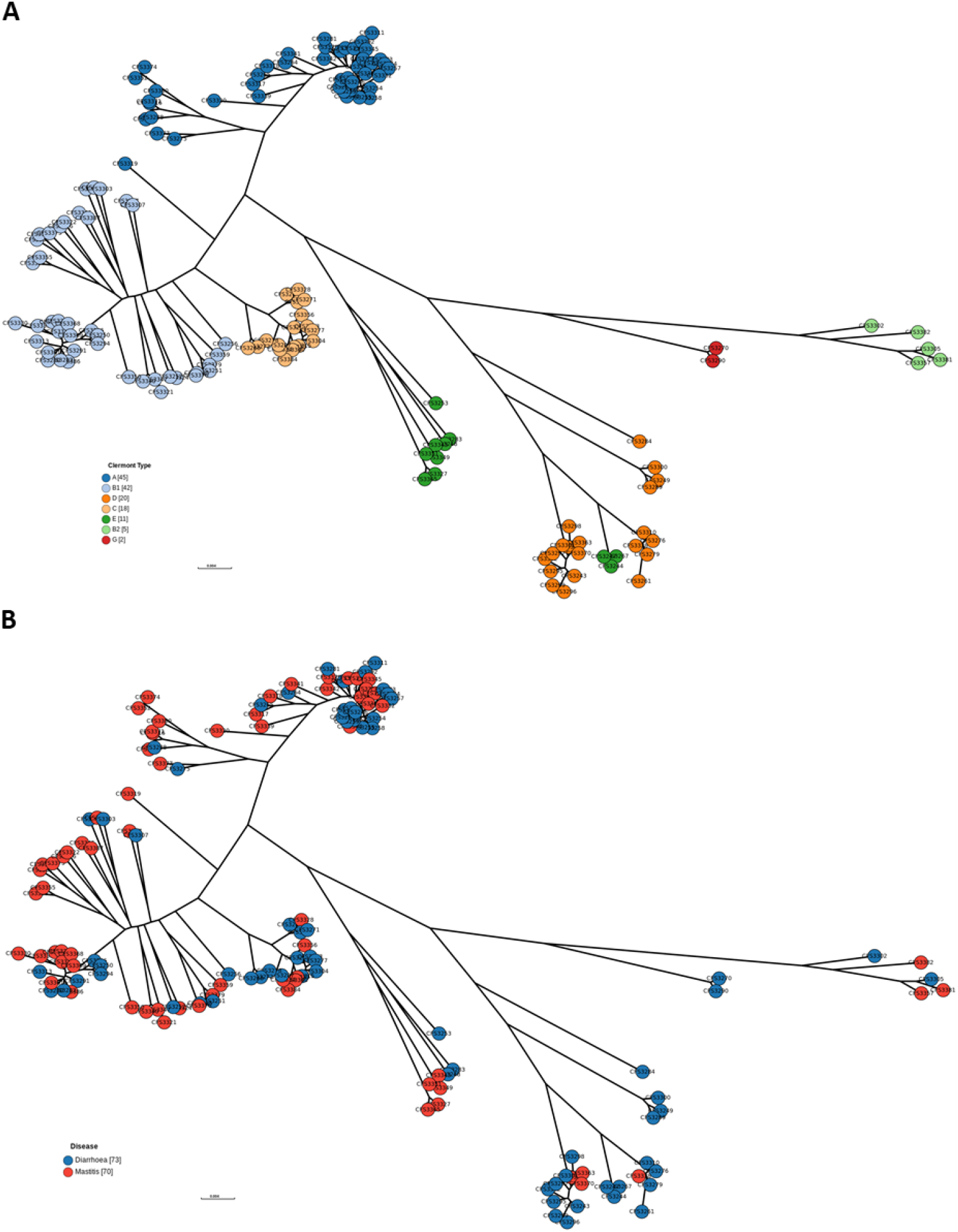
SNP tree generated in Enterobase, using CFS3246 as the reference, and visualised using GrapeTree. *E. coli* isolates were coloured in based on the Clermont Type phylogrouping scheme **(A)** and disease that the sample was recovered from **(B)**.

### AMR genotypes identified by database comparisons

AMR genes were identified using AMRFinder and ResFinder databases. All ResFinder database genes were also detected by AMRFinder, with AMRFinder providing additional genes. All isolates encoded multidrug efflux RND transporter permease subunit gene *acrF* and a version of *bla*EC gene encoding beta-lactam resistance, and 134 isolates encoded multidrug efflux MFS transporter *mdtM* (**Figure 2**). When the efflux pump genes were excluded, 82 isolates encoded more than 1 AMR gene responsible for resistance to a specific antibiotic. The most common gene identified was *bla*_TEM-1_ which is associated with beta-lactam resistance, being detected in 53 isolates (37.06%) (**Table 2**). This gene confers resistance to cephalothin and amoxicillin. Other genes of high frequency include *tet*(A) (30.77%), and *sul2* (35.66%) which are associated with tetracycline and sulfonamide resistance, respectively. There was a high prevalence of resistance to tetracycline: five different genes conferred resistance to tetracycline with *tet*(A) and *tet*(B) (16.08 %) being the most prevalent. Trimethoprim-sulfamethoxazole resistance requires the presence of genes from both *sul* and *dfr* gene groups. There was relatively low resistance detected for the aminoglycoside antibiotic gentamicin with 15 of the 143 genomes harbouring gentamicin resistance-encoding genes. The most frequent AMR gene promoting gentamicin resistance was *ant(2’’)-Ia*, which was detected in 8 of the 143 isolates. Florfenicol resistance marked by the presence of the *floR* gene was detected in 14 isolates (9.79%).

**Table 2:**
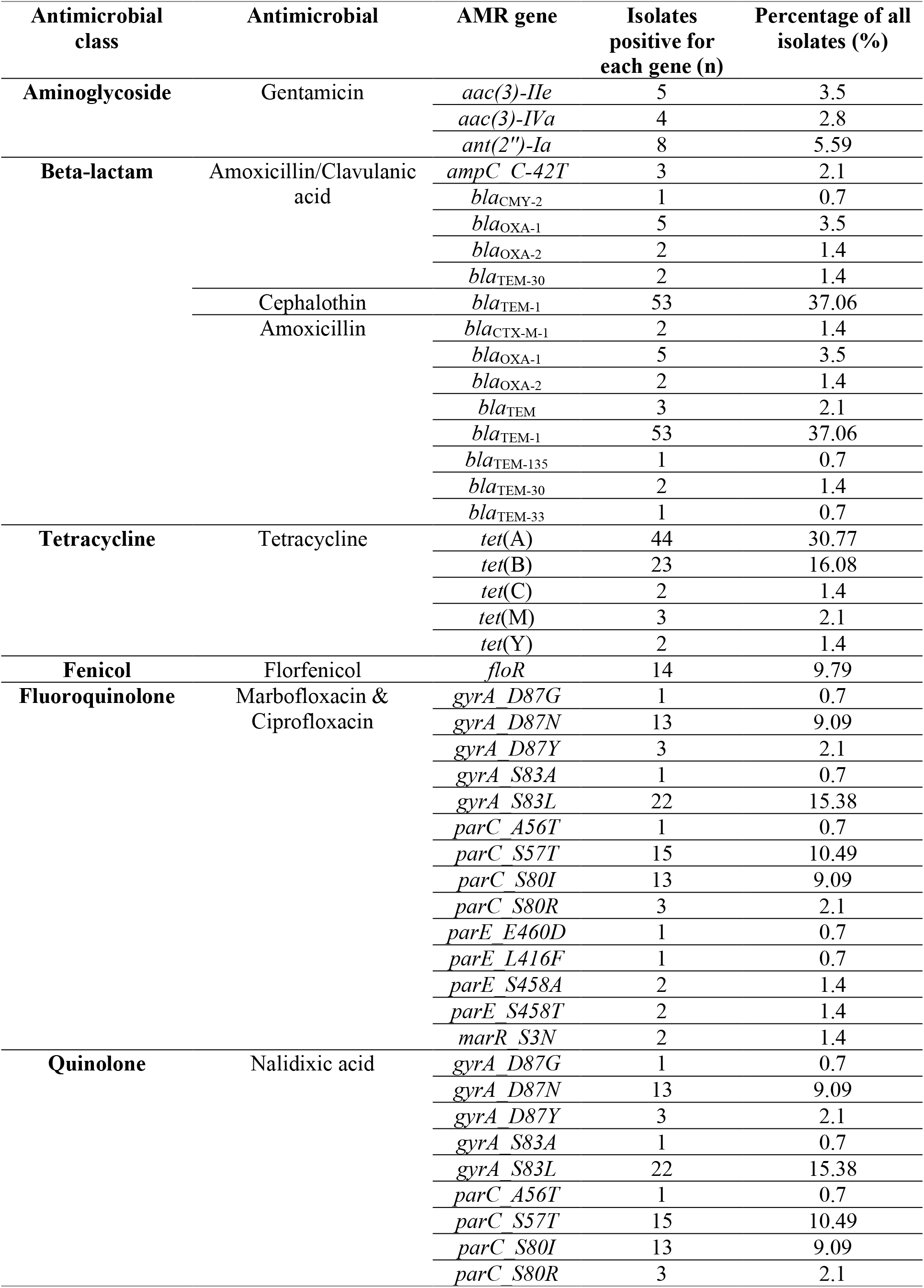
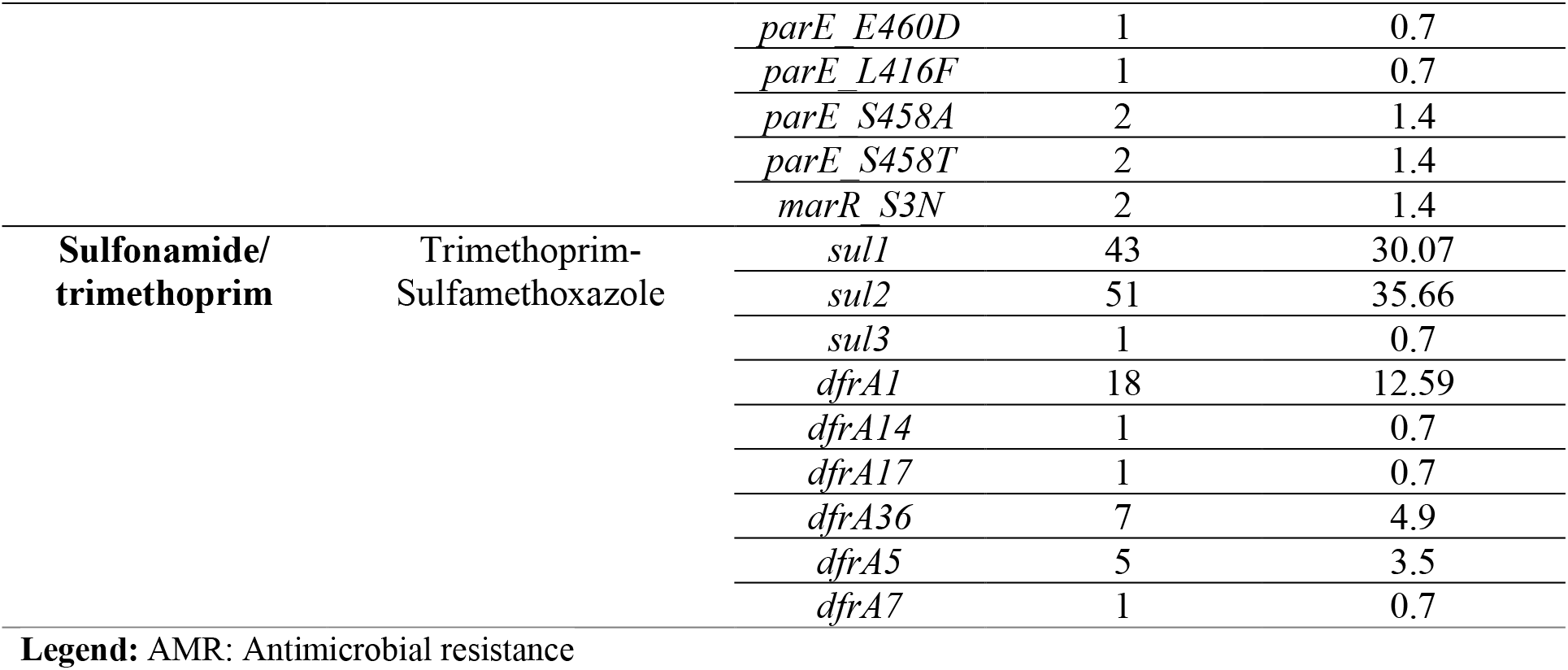
Antimicrobial resistance genes identified in *Escherichia coli* isolates of bovine origin isolated from animals suffering from diarrhoea and mastitis (n=143)

**Figure 2:**
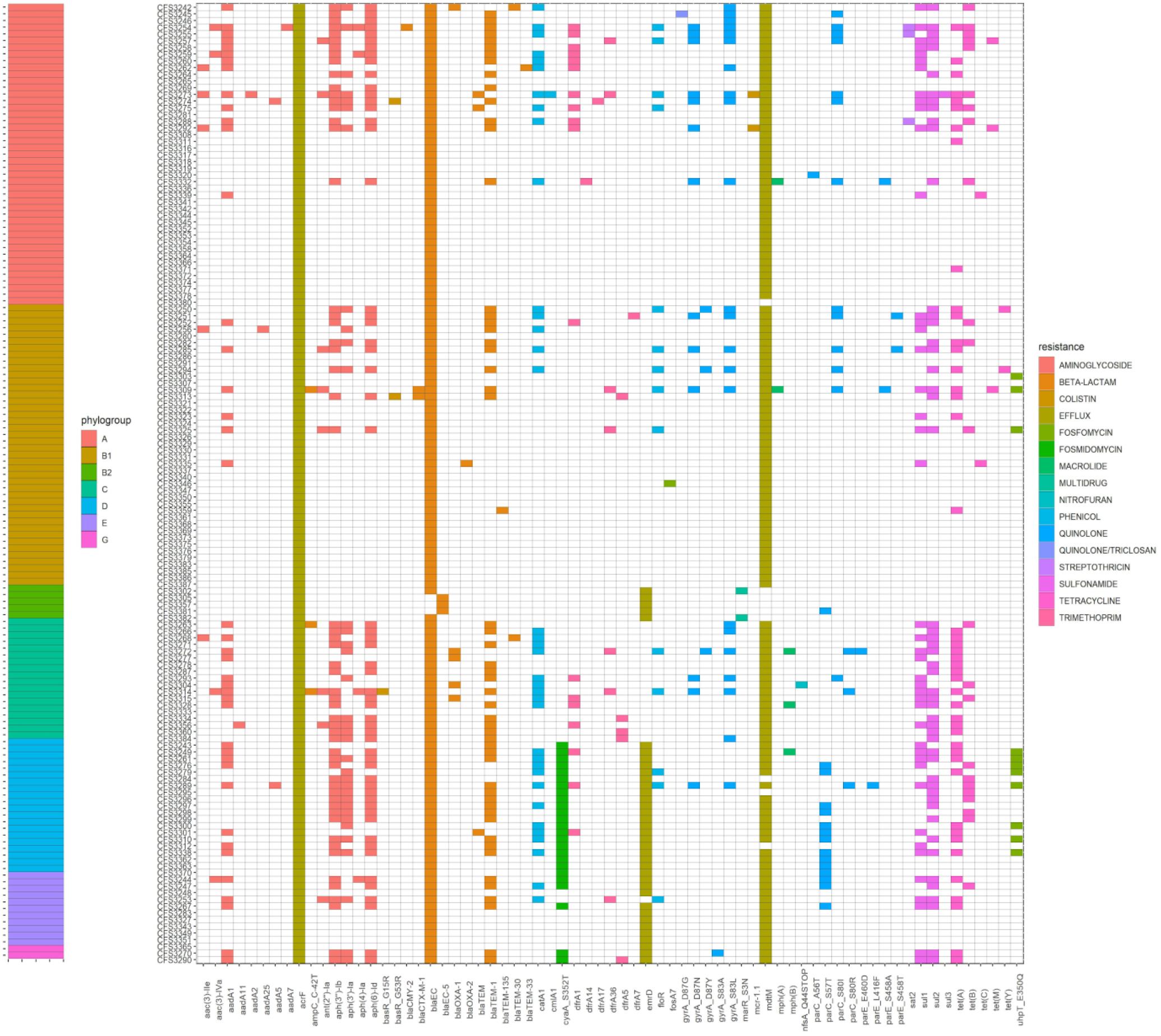
A heatmap showing an absence/presence matrix detailing AMR genotypes and associated phylogroups for 143 *Escherichia coli* isolates of bovine origin. A colour key is shown to the right indicating the classes of compounds tested.

There was a high level of antimicrobial resistance present in the cohort of isolates tested, 51.04% (n=73) of which had an MDR genotype harbouring resistance to between 3 and 10 of the antibiotics investigated for the correlation in this study. The antimicrobial therapy options against Gram-negative MDR bacteria are limited. This poses a risk to veterinary public health and a challenge in the fight against nosocomial infections (Giamarellou and Poulakou, 2009). The AMR genes most frequently detected in the isolates (*bla*_TEM-1_, *tet*(A) and *sul2*) agreed with previous studies investigating AMR genes in MDR *E. coli* of bovine origin, which detailed a high level of tetracycline, sulfonamide and beta-lactam resistance among isolates sampled from cattle in different European countries (Karczmarczyk et al., 2011).

### AST by disk diffusion (AST-DD)

A total of 10 antibiotics were investigated by conventional disk diffusion testing for correlation with genotype determination. These compounds belonged to 7 classes of antimicrobial agent which included aminoglycoside, beta-lactam, tetracycline, phenicol, fluoroquinolone, quinolone and sulfonamide-trimethoprim compounds. Of the 143 isolates tested, 46 (32.17%) *E. coli* were fully susceptible to all 10 antibiotics tested for by disk diffusion. All other isolates were resistant or intermediately resistant to between 1 and 10 antibiotics. AST-DD determined that 45.5 % (65/143) of isolates were multi-drug resistant (MDR). The most frequent resistance recorded was seen for the antibiotics amoxicillin and tetracycline with 66 and 65 resistant isolates (46% and 45%), respectively (**Figure 3A**). The antibiotics demonstrating least resistance among isolates recorded were cephalothin (7.69%) and amoxicillin/clavulanic acid (8.39%).

**Figure 3:**
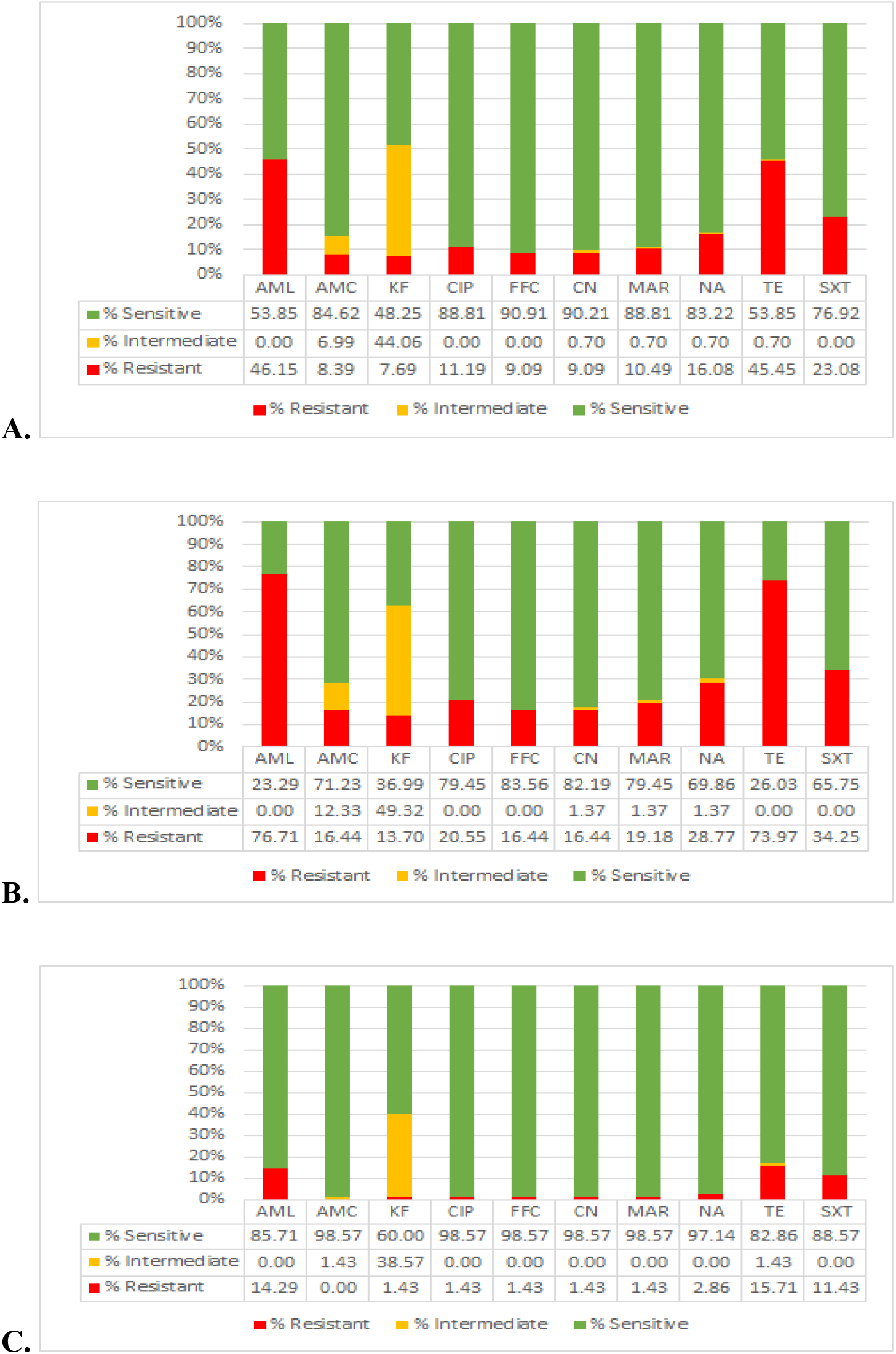
A bar graph showing the disk diffusion susceptibility profile for all isolates (**A**; n = 143), isolates of diarrhoeal origin (**B**; n = 73) and mastitis origin (**C**; n = 70). AML – Amoxicillin, AMC – Amoxicillin/Clavulanic acid, KF – Cephalothin, CIP – Ciprofloxacin, FFC – Florfenicol, CN – Gentamicin, MAR – Marbofloxacin, NA – Nalidixic acid, TE – Tetracycline, SXT – Trimethoprim-Sulfamethoxazole.

Resistance patterns of the isolates were compared based on disease type (diarrhoea, **Figure 3B***versus* mastitis, **Figure 3C**). There was a higher prevalence of antimicrobial resistance in the cohort of diarrhoeal samples in comparison to mastitis samples. The highest percentage resistance recorded for diarrhoeal samples was 76.7 % for amoxicillin, while for mastitis isolates it was 15.7 % for tetracycline. No resistance to amoxicillin/clavulanic acid was recorded for mastitis samples, and low resistance to ciprofloxacin, florfenicol, gentamicin, marbofloxacin and nalidixic acid (1.4 %) was detected. Overall, there was a higher prevalence of resistant isolates among samples recovered from cases of diarrhoea.

### AST by minimum inhibitory concentration (AST-MIC)

A subset of 82 isolates with more than one AMR-encoding gene was selected for MIC determination. Of these isolates, 23.17 % were sourced from mastitis cases, while 76.83 % were sourced from cases of diarrhoea. A total of 6 antibiotics were investigated for correlation between both methods. These antibiotics belonged to 5 classes of antimicrobial agent. Of the 82 isolates tested, 11 (13.41 %) *E. coli* were fully susceptible to all 6 compounds tested. All other isolates were resistant or intermediately resistant to between 1 and 6 antibiotics. The most frequent resistance was recorded for the antimicrobial compounds tetracycline and trimethoprim-sulfamethoxazole with 60 and 33 resistant isolates (73.17 % and 40.24 %), respectively (**Figure 4**). The antibiotics showing least resistant isolates recorded were cephalothin (12.20 %) and gentamicin (14.63 %). As with AST-DD, the antimicrobial resistance was higher among those isolates recovered from cases of diarrhoea (n = 63), rather than mastitis (n = 19).

**Figure 4:**
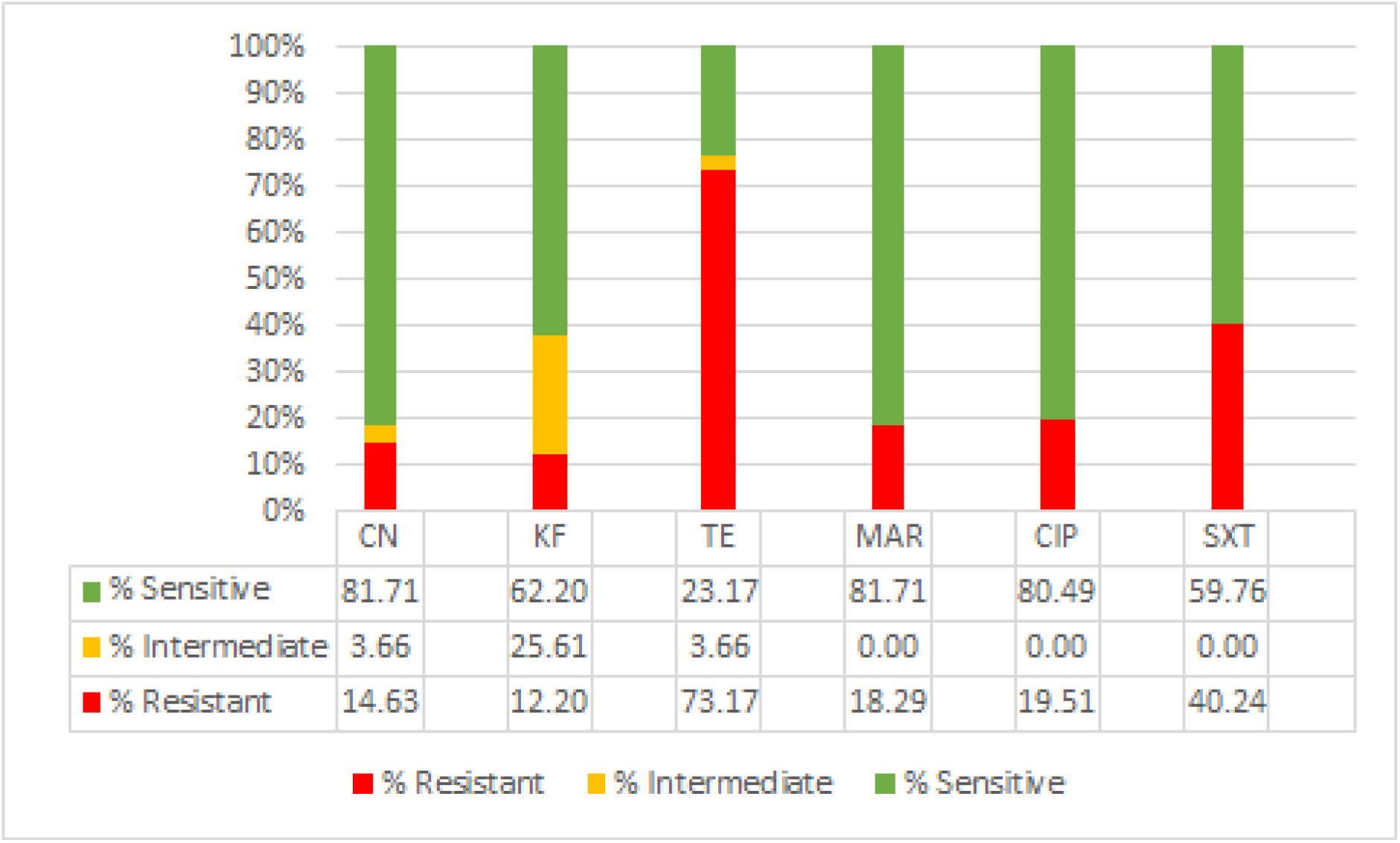
MIC susceptibility profile for all isolates (n=82). CN – Gentamicin, KF – Cephalothin, TE – Tetracycline, MAR – Marbofloxacin, CIP – Ciprofloxacin and SXT – Trimethoprim-Sulfamethoxazole.

As with AST-DD, when comparing resistance profiles based on disease type for AST-MIC, there was a higher prevalence of antimicrobial resistance in the cohort of diarrhoeal samples in comparison to mastitis samples. The highest percentage resistance recorded for diarrhoeal samples was 79.37 % for tetracycline while for mastitis isolates it was 52.63 % for tetracycline. No resistance to gentamicin was recorded for mastitis samples while low resistance to ciprofloxacin (5.26 %), marbofloxacin (5.26 %) and cephalothin (5.26 %) was noted. Overall, there was a higher prevalence of resistant isolates among diarrhoeal samples. Overall, both when determined by AST-DD and AST-MIC, the resistance phenotypes of study isolates tended to differ according to the disease they were associated with. Interestingly, the isolates did not cluster on the phylogenetic tree according to disease type, with mastitis and diarrhoeal samples spread evenly across phylogroups and wgMLST types. This suggests that the difference in resistance phenotype could be due to a selective pressure due to the antibiotics used to treat these infections. Mastitis is predominantly treated with amoxicillin and other beta-lactam antibiotics which correlates well with the resistance detected in mastitis isolates. Broader spectrum antibiotics such as tetracycline are usually prescribed to treat bovine diarrhoea (Smith, 2015). However, digestive pathogens also tend to be more exposed to antibiotics than mastitis pathogens, which could also explain the higher rate of resistance.

### Correlation of genotypic and phenotypic methods (Disk Diffusion)

Correlation between genotypic and phenotypic data in this study was assessed by means of a two-by-two table analysis. Correlation of AST-WGS with AST-DD for all antibiotics overall was 92.5% for test specificity and 98.8% for test sensitivity (**Table 3**). The kappa score (κ = 0.81) indicates almost perfect agreement. Correlation for individual antibiotics varied with five compounds yielding almost perfect agreement (κ > 0.90), one showing substantial agreement and four showing moderate agreement.

**Table 3:** A table showing the statistical correlation between genotypic (based on whole genome sequencing) and phenotypic (as determined by disk diffusion) methods of AMR detection, test performances and kappa correlations of *Escherichia coli* isolates of bovine origin (n=143).

| Antibiotic | P+/G+ | P-/G- | G+/P- | G-/P+ | Specificity | Sensitivity | PPV | NPV | $\kappa$ |
| --- | --- | --- | --- | --- | --- | --- | --- | --- | --- |
| **Amoxicillin** | 65 | 77 | 0 | 1 | 100 | 98.5 | 100 | 98.7 | 0.99 |
| **Amoxicillin/**<br>**Clavulanic acid** | 10 | 120 | 1 | 12 | 99.2 | 45.5 | 90.9 | 90.9 | 0.56 |
| **Cephalothin** | 42 | 58 | 11 | 32 | 84.1 | 56.8 | 79.2 | 64.4 | 0.4 |
| **Ciprofloxacin** | 16 | 101 | 26 | 0 | 79.5 | 100 | 38.1 | 100 | 0.47 |
| **Florfenicol** | 13 | 129 | 1 | 0 | 99.2 | 100 | 92.9 | 100 | 0.96 |
| **Gentamicin** | 14 | 128 | 1 | 0 | 99.2 | 100 | 93.3 | 100 | 0.96 |
| **Marbofloxacin** | 16 | 101 | 26 | 0 | 79.5 | 100 | 38.1 | 100 | 0.47 |
| **Nalidixic acid** | 24 | 101 | 18 | 0 | 84.9 | 100 | 57.1 | 100 | 0.65 |
| **Tetracycline** | 64 | 76 | 1 | 2 | 98.7 | 97 | 98.5 | 97.4 | 0.96 |
| **Trimethoprim-**<br>**Sulfamethoxazole** | 32 | 110 | 0 | 1 | 100 | 97 | 100 | 99.1 | 0.98 |
| **Overall** | 150 | 516 | 46 | 3 | 92.5 | 98.8 | 77.4 | 99.3 | 0.81 |

**Kappa correlation ( $\kappa$ ):**0.01–0.20 (none to slight agreement), 0.21–0.40 (fair agreement), 0.41– 0.60 (moderate agreement), 0.61–0.80 (substantial agreement), 0.81–1.00 (almost perfect agreement).

Based on these calculations, an almost perfect agreement was observed between AST-WGS and AST-DD for the antibiotics florfenicol, tetracycline, gentamicin, trimethoprim-sulfamethoxazole and amoxicillin. These had a κ > 0.95 and all had a correlation specificity > 98.7% and sensitivity > 97.0%, which suggests AST-WGS genotype calling could be used as an accurate predictor of AMR phenotype for these compounds. Both amoxicillin and trimethoprim-sulfamethoxazole had a test specificity of 100% with no false negative results. A large number of genotypes conferred resistance to amoxicillin (**Figure 5A**), all of which can be considered good predictors of resistance due to the absence of false positive results. There was also an almost perfect agreement for the antibiotic florfenicol (**Figure 6A**). Only one genotype (*floR*) conferred resistance to florfenicol and there was a clear distinction between resistant and susceptible isolates. The gene that seems to most reliably predict resistance to gentamicin (**Figure 6B**) is *aac(3)-IIe*. The least reliable gene seems to be *ant(2’’)-Ia* as it was found to be present in a susceptible isolate. The prediction of tetracycline resistance by AST-WGS also had an almost perfect agreement between the methods. There were a large number of genotypes that confer resistance and susceptible isolates that had no associated genotype (**Figure 6C**). In the case of trimethoprim-sulfamethoxazole both *dfr* and *sul* genes were required to be present to confer resistance (**Figure 6D**). There were no false positive results recorded for this antibiotic combination and differences between resistant and susceptible were clear due to the absence of intermediately resistant isolates.

**Figure 5:**
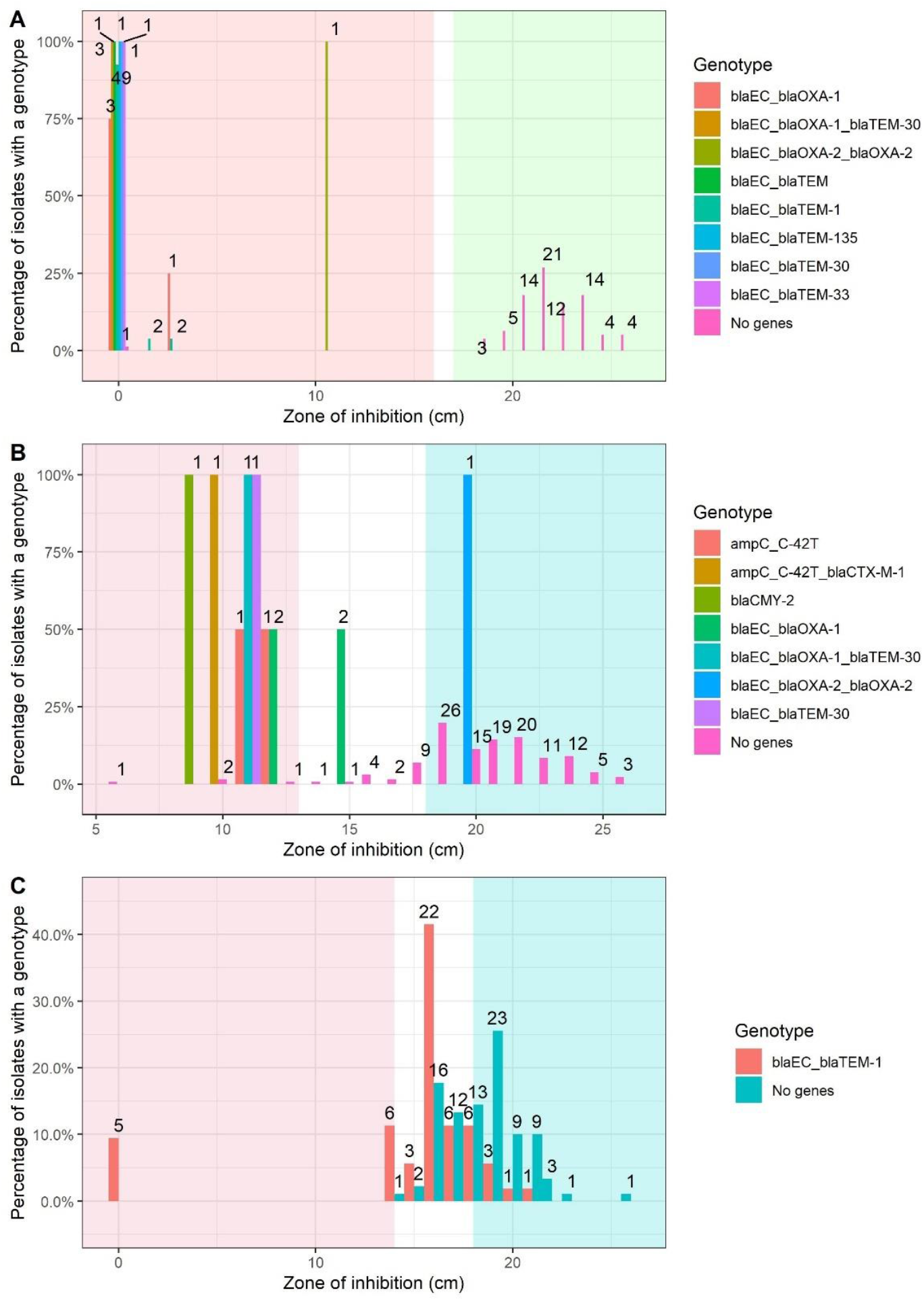
Bar charts showing the distribution of isolates with specific resistant genotypes at each zone of inhibition for beta-lactam antibiotics: amoxicillin **(A),** amoxicillin-clavulanic acid **(B),** cephalothin **(C).** Numbers shown at the top of each column represents the number of isolates with the specific genotype, at that zone of inhibition. Zones of inhibition represent an average of 3 independent measurements and were rounded up to integer level. Red shaded background denotes the zone of inhibition for which the isolates are considered to be resistant, while blue/green shaded background denotes the zone of inhibition for which the isolates are considered to be susceptible. White background represents zones of inhibition of intermediately resistant isolates.

**Figure 6:**
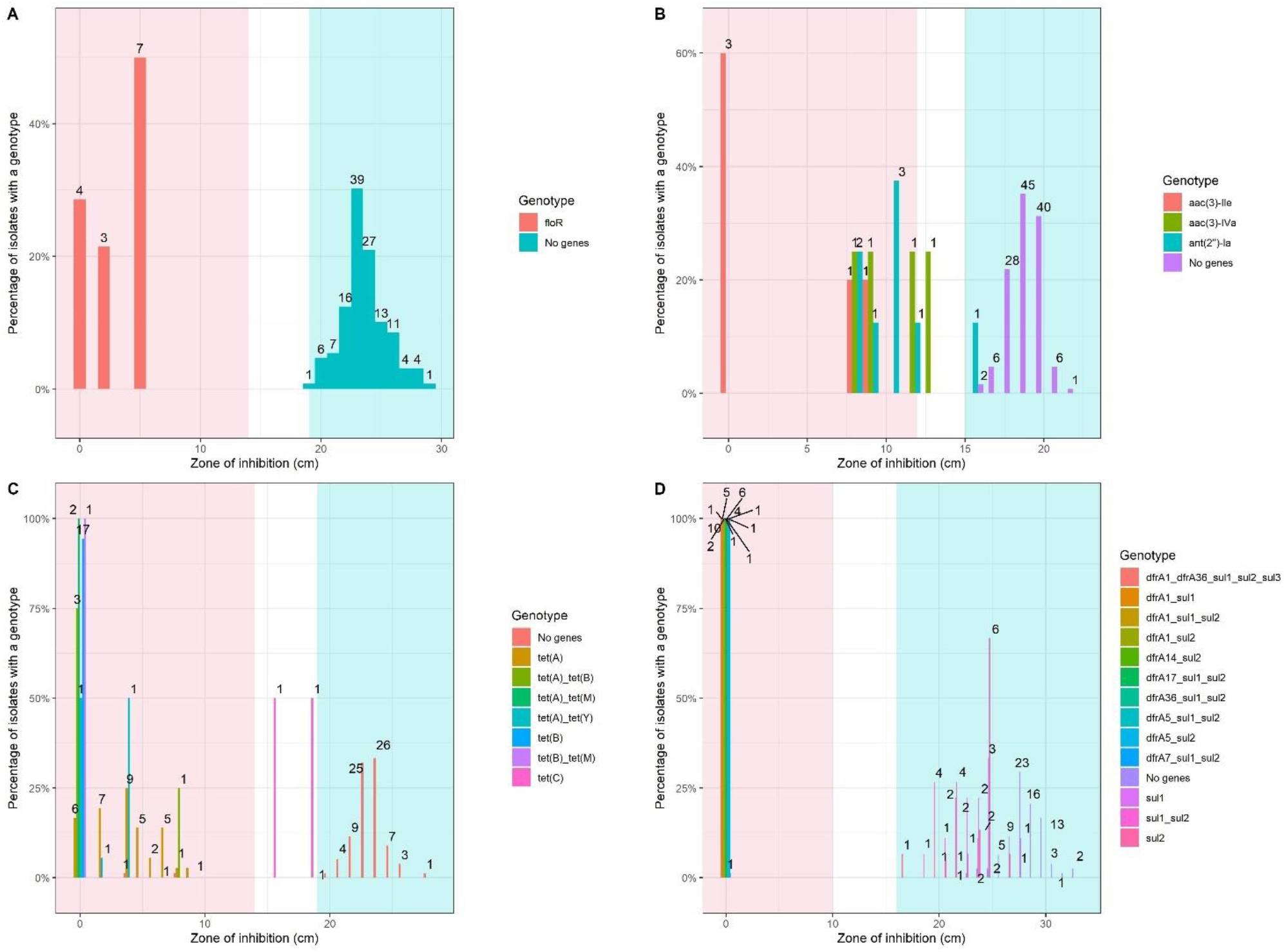
Bar charts showing the distribution of isolates with specific resistant genotypes at each zone of inhibition for florfenicol **(A)**, gentamicin **(B),** tetracycline **(C)** and trimethoprim-sulfamethoxazole **(D).** Number at each column represents the number of isolates with the specific genotype, at that zone of inhibition. Zones of inhibition represent an average of 3 independent measurements and were rounded up to integer level. Red background denotes the zone of inhibition for which the isolates are considered to be resistant, while blue/green background denotes the zone of inhibition for which the isolates are considered to be susceptible. White background represents zones of inhibition of intermediately resistant isolates.

Substantial agreement (κ > 0.6) was observed for nalidixic acid with a κ value of 0.65. Similarly to the antimicrobial agents discussed previously, nalidixic acid had no false negative results, giving the compound 100% NPV in the correlation. The correlation for this antibiotic could be improved by investigating the genes causing false positive results for 18 of the isolates (**Figure 7B**).

**Figure 7:**
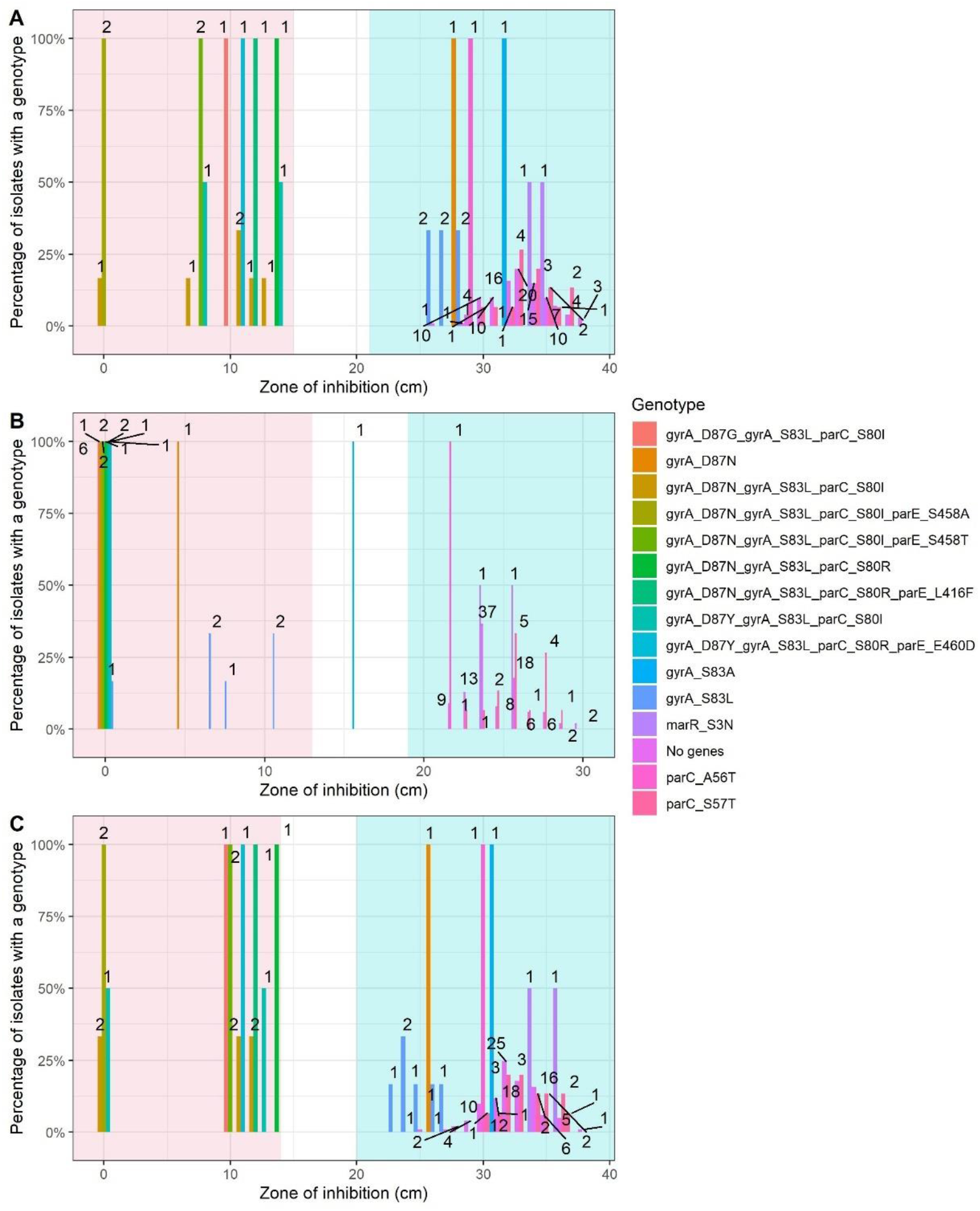
Bar charts showing the distribution of isolates with specific resistant genotypes at each zone of inhibition for quinolones: ciprofloxacin **(A),** nalidixic acid **(B)** and marbofloxacin **(C).** Number at each column represents the number of isolates with the specific genotype, at that zone of inhibition. Zones of inhibition represent an average of 3 independent measurements and were rounded up to integer level. Red background denotes the zone of inhibition for which the isolates are considered to be resistant, while blue/green background denotes the zone of inhibition for which the isolates are considered to be susceptible. White background represents zones of inhibition of intermediately resistant isolates.

Moderate agreement was observed for amoxicillin/clavulanic acid, cephalothin, ciprofloxacin and marbofloxacin. In the case of amoxicillin/clavulanic acid, which had a test specificity of 99.2%, test sensitivity (45.5 %) negatively impacted correlation resulting in a Cohen’s kappa score of 0.56. For amoxicillin/clavulanic acid, *bla*_OXA-2_ was not an accurate predictor of resistance resulting in a false positive (**Figure 5B**). There were also a large number of false negative results for this antibiotic which suggests there are genotypes conferring amoxicillin/clavulanic acid resistance that are undiscovered or not included in the AMRFinder database. For the correlation with cephalothin, high levels of false positive and false negative results were recorded, as 11 susceptible isolates with AMR genotypes were present and 32 isolates were intermediately resistant with others being recorded as resistant with no AMR genotype respectively (**Figure 5C**). It is therefore apparent that further consideration should be given to those genes that confer resistance to beta-lactam antibiotics due to the presence of resistant isolates with no genotype. The fluoroquinolones ciprofloxacin and marbofloxacin also were moderately correlated, both with a κ value of 0.47. Both antimicrobial agents also had the same test performances in terms of test specificity (79.5%) and test sensitivity (100.0%). There were no false negative results for these antibiotics but the poorer correlation is likely due to the high number of false positive results i.e., isolates that are phenotypically susceptible but have an associated genotype (**Figure 7**, isolates in the blue/green shaded area with an associated genotype). Genes that were poor predictors of AMR in this class of antibiotic, causing false positive results in the correlation, included *gyrA_S83L* and *parC_S57T* (**Figure 7A & 7C**). Mutations in the gene encoding the A subunit of DNA gyrase (*gyrA)* and topoisomerase IV subunit ParC gene (*parC)* are the major cause of quinolone resistance encountered in clinical isolates. Studies have revealed that these mutations predominate in conserved regions known as the quinolone-resistance-determining region or QRDR, involved in DNA binding. In GyrA, the QRDR comprise amino acids between 67 and 106 (Varughese et al., 2018). The most common mutations discovered in gram-negative bacteria such as *E*.*coli*, *Shigella*, *Citrobacter* and *Pseudomonas* were at codon 83 and 87 in GyrA (Ruiz, 2003), which were the only mutations found in this study. In *E. coli*, *gyrA* mutations usually occur at serine-83, which is substituted by leucine or tryptophan and causes high resistance to quinolones. However, the replacement of serine by alanine causes a lower resistance level (Ruiz, 2003; Krishnan et al., 2012). In our study, the *gyrA_S83A* gene resulted in a strain which was phenotypically susceptible to ciprofloxacin and marbofloxacin and intermediately resistant to nalidixic acid (**Figure 7**), therefore confirming the findings of previous studies. Mutations in *parC* usually occur at 78, 80 and 84 positions (Hiasa, 2002; Ito et al., 2008), which calls into question the point mutations we observed in positions 56 and 57, with all strains harbouring *parC_A56T* and *parC_S57T* susceptible to all 3 quinolones in the study. Finally, studies show that multiple mutations are required for high level of resistance (Fàbrega et al., 2008; Shigemura et al., 2012), which is also consistent with our findings.

Most of the discrepancies in correlation were due to false negative results, i.e., isolates that were phenotypically resistant but genotypically susceptible. This is most likely due to the presence of new AMR gene variants that have not yet been described. False negatives may also be caused by other resistance mechanisms such as the action of efflux pumps (Stubberfield et al., 2019). Two efflux pump genes, *acrF* and *mdtM*, which were identified in all study isolates, may partially explain the discrepancy between the 81 genotypically resistant isolates and the 135 phenotypically resistant isolates. In *Escherichia coli*, the AcrAB-TolC efflux pump plays an intrinsic role in resistance to hydrophobic solvents. The *acrEF* operon encodes the components of a second efflux pump with AcrE and AcrF highly homologous to AcrA and AcrB, and can confer the same solvent resistance (Kobayashi et al., 2001). However, both *acrE* and *acrF* would be necessary for the resistance mechanism, making the role of *acrF* only in the resistance of the isolates in our study unclear. The multidrug resistance protein MdtM, a recently characterised homologue of MdfA and a member of the major facilitator superfamily, functions in alkaline pH homeostasis (Holdsworth and Law, 2013). Both *acrEF* and *mdtM* (also known as *yjiO*) have been found to confer resistance to some antibiotics as well as other compounds such as ethidium bromide (Nishino and Yamaguchi, 2001).

False positives i.e., isolates that are genotypically resistant and phenotypically susceptible to certain antimicrobial compounds, account for the remaining discrepancies. False positives may be caused by detection of silenced genes that do not express resistance (Enne et al., 2006). Genotypically resistant, phenotypically susceptible isolates may also arise if the wrong AMR genes are selected to correlate genotype to phenotype. This highlights the need for careful curation of AMR gene alleles conferring resistance to each antimicrobial compound in order to ensure reliable and accurate prediction of AMR.

Poor correlation between genotype and phenotype was often accompanied by a high proportion of intermediately resistant isolates (**Table 4**). Higher percentages of intermediately resistant strains were found for antibiotics with κ < 0.6. Notably, 44.1% isolates were intermediately resistant to cephalothin which had a κ correlation of 0.40. Cephalothin had the highest proportion of intermediately resistant isolates in this study, and was the likely cause of the poor correlation. The high number of isolates which were intermediately resistant to cephalothin may be due to issues with the method. Previous studies comparing reference broth dilution MIC and disk diffusion methods for cephalothin AST determination highlighted discrepancies in results (Nguyen and Graber, 2013). This discrepancy was also highlighted in our study. Of the 63 isolates which were intermediately resistant to cephalothin by disk diffusion, 44 were captured in the sub-selection of 82 isolates tested by AST- MIC. Only 21 of this sub-selection were then intermediately resistant when tested by AST-MIC.

**Table 4:** Number and percentage of intermediately resistant strains for each antibiotic

| Antibiotic | No. of<br>intermediately<br>resistant strains | % of Total<br>isolates | $\kappa$ |
| --- | --- | --- | --- |
| Amoxicillin | 0 | 0 | 0.99 |
| Amoxicillin/ Clavulanic acid | 10 | 7 | 0.56 |
| Cephalothin | 63 | 44.1 | 0.4 |
| Ciprofloxacin | 0 | 0 | 0.47 |
| Florfenicol | 0 | 0 | 0.96 |
| Gentamicin | 1 | 0.7 | 0.96 |
| Marbofloxacin | 1 | 0.7 | 0.47 |
| Nalidixic acid | 1 | 0.7 | 0.65 |
| Tetracycline | 1 | 0.7 | 0.96 |
| Trimethoprim-Sulfamethoxazole | 0 | 0 | 0.98 |

### Correlation of genotypic and phenotypic methods (MIC)

Correlation of AST-WGS with AST-MIC for all antibiotics overall was 75 % for test specificity and 95.9 % for test sensitivity (**Table 5**). Two antibiotics had a κ > 0.90 which indicated a near perfect agreement between the methods, and notably, the κ correlation for trimethoprim-sulfamethoxazole was 1, with no false positives or negatives for this antibiotic. Fair agreement was observed for some compounds such as cephalothin, ciprofloxacin and marbofloxacin.

**Table 5:**
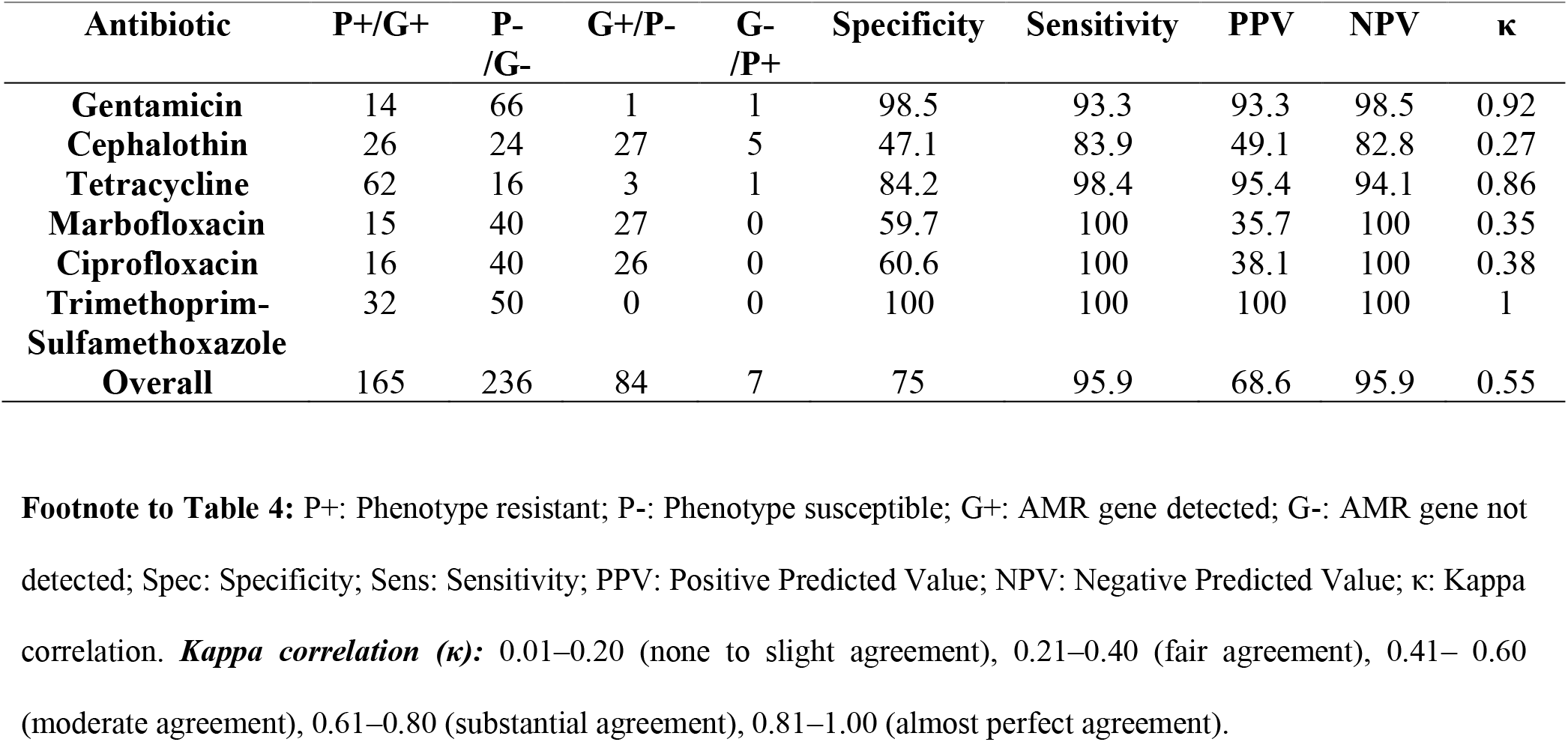
Correlation between genotypic (whole genome sequencing) and phenotypic (MIC) methods of AMR detection, test performances and kappa correlations of *Escherichia coli* isolates of bovine origin (n=82)

| Antibiotic | P+/G+ | P-<br>/G- | G+/P- | G-<br>/P+ | Specificity | Sensitivity | PPV | NPV | κ |
| --- | --- | --- | --- | --- | --- | --- | --- | --- | --- |
| **Gentamicin** | 14 | 66 | 1 | 1 | 98.5 | 93.3 | 93.3 | 98.5 | 0.92 |
| **Cephalothin** | 26 | 24 | 27 | 5 | 47.1 | 83.9 | 49.1 | 82.8 | 0.27 |
| **Tetracycline** | 62 | 16 | 3 | 1 | 84.2 | 98.4 | 95.4 | 94.1 | 0.86 |
| **Marbofloxacin** | 15 | 40 | 27 | 0 | 59.7 | 100 | 35.7 | 100 | 0.35 |
| **Ciprofloxacin** | 16 | 40 | 26 | 0 | 60.6 | 100 | 38.1 | 100 | 0.38 |
| **Trimethoprim-Sulfamethoxazole** | 32 | 50 | 0 | 0 | 100 | 100 | 100 | 100 | 1 |
| **Overall** | 165 | 236 | 84 | 7 | 75 | 95.9 | 68.6 | 95.9 | 0.55 |

Almost perfect agreement was observed between AST-WGS and AST-MIC for the antibiotics gentamicin, tetracycline and trimethoprim-sulfamethoxazole. These had κ > 0.80 values and a correlation specificity > 84 % and sensitivity > 90 %, which suggests AST-WGS could be used as an accurate predictor of AMR for these antibiotics. For gentamicin (**Figure 8B**), the gene that seems to most reliably predict resistance is *aac(3)-IIe*. The least reliable gene seems to be *ant(2’’)-Ia* as it is present in a susceptible isolate. This finding agrees with the results for gentamicin AST-DD. Tetracycline resistance was predicted well by AST-WGS, with almost perfect agreement between the methods. There were a large number of genotypes that confer resistance and susceptible isolates that had no associated genotype (**Figure 8C**). Once again, both *dfr* and *sul* genes were both required to confer resistance to trimethoprim-sulfamethoxazole (**Figure 8D**). Differences between resistant and susceptible were clear due to the absence of isolates intermediately resistant to trimethoprim-sulfamethoxazole.

**Figure 8:**
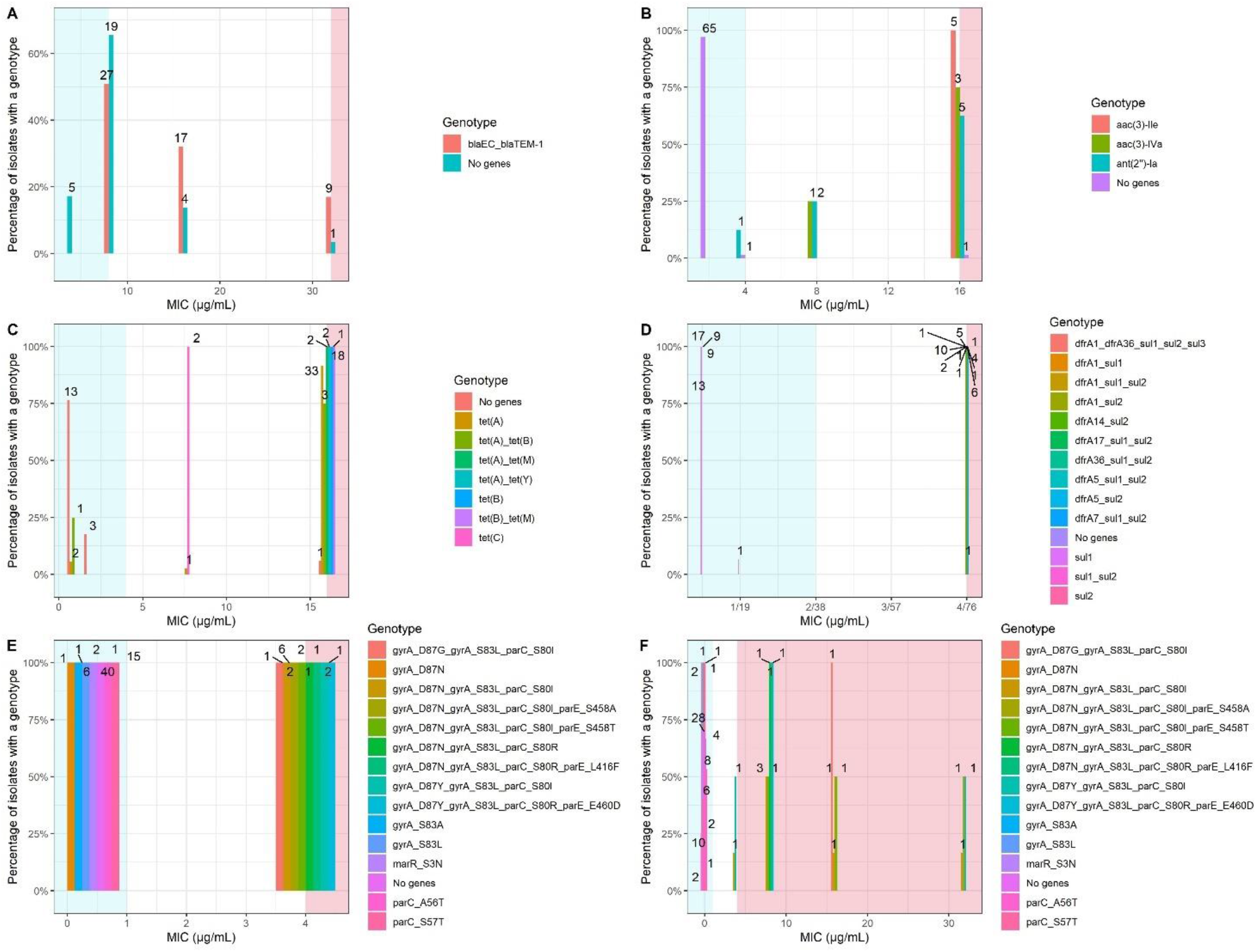
Bar charts showing the distribution of isolates with specific resistant genotypes at each MIC for cephalothin **(A),** gentamicin **(B),** tetracycline **(C),** trimethoprim-sulfamethoxazole **(D),** ciprofloxacin **(E)** and marbofloxacin **(F).** Number at each column represents the number of isolates with the specific genotype, at that MIC. Red background denotes the MIC for which the isolates are considered to be resistant, while blue/green background denotes the MIC for which the isolates are considered to be susceptible. White background represents MIC of intermediately resistant isolates.

For cephalothin, there was an increase in false positive results and a decrease in false negative results in comparison to AST-DD (**Figure 8A**). A total of 27 susceptible isolates with AMR genotypes were present (*versus* 11 as determined by AST-DD) and 5 isolates were intermediately resistant without an AMR genotype respectively (*versus* 32 as determined by AST-DD). A better appreciation of the linkages between genotypes and phenotypes that confer resistance to beta-lactam antibiotics is required.

Fluoroquinolones ciprofloxacin and marbofloxacin returned a fair agreement between the methods also, with a κ value of 0.38 and 0.35 respectively. Both antimicrobial compounds also had the same test performances in terms of test sensitivity (100.0 %) but differed in terms of test specificity. No false negative results were recorded for these compounds; therefore, the poorer correlation is likely owed to the high number of false positive results (**Figures 8E & 8F,** isolates in the blue/green shaded area with an associated genotype).

### General discussion – comparison between disk diffusion and MIC correlations

In this study, WGS was explored as an alternative means of predicting AMR in place of conventional disk diffusion and MIC determined by microbroth dilution, which are routinely used phenotypic methods of AST. WGS was carried out on 143 *Escherichia coli* of bovine origin and AMR genes detected using bioinformatic analysis. The same isolates were then subjected to AST-DD for a total of 10 antibiotics of clinical and veterinary importance. A subset of these same isolates were also subjected to AST-MIC analysis using Sensititre plates. Genotypic and phenotypic AMR results were correlated and statistical agreement between the methods was determined.

Previous studies reported on the correlation between AST-MIC and genotypic AMR patterns (Card et al., 2013; Stoesser et al., 2013; Zankari et al., 2013; Tyson et al., 2015; Feldgarden et al., 2019; Stubberfield et al., 2019) but no study to date examined the correlation between AST-DD and genotypic AMR patterns. While broth microdilution is the clinical standard for AST, disk diffusion is a cheaper, and a more accessible alternative. AST-DD correlated better with AST-WGS than AST-MIC. The overall κ value for correlation of AST-DD with AST-WGS was 0.81, in comparison to 0.55 for AST-MIC by Sensititre with AST-WGS. This study highlights AST-DD is equivalently as useful a method of AST as AST-MIC.

The accuracy of genomic methods such as AST-WGS was determined by the reliability of the software that is used to determine AMR. The ResFinder and AMRFinder databases which were used to predict AMR genes, as previously discussed, may not always correctly identify AMR genes. Previous studies implementing ResFinder have highlighted this feature (Feldgarden et al., 2019). An isolate which is wrongly classified as genotypically susceptible due to lack of known genetic determinant of resistance may negatively affect treatment decisions. Although AST-WGS can offer more information on the test isolates, a disadvantage is that genomic approaches cannot always predict a phenotype, as level of gene expression and protein production from identified genes may differ between strains. Bacteria have gene silencing mechanisms, and mutations may create stop codons in the data (Dryselius et al., 2003).

Similar studies that correlated WGS with phenotypic AST found that WGS could be used as an accurate predictor of AMR in human, bovine or porcine isolates (Zankari et al., 2012; Card et al., 2013; Stoesser et al., 2013; Tyson et al., 2015; Feldgarden et al., 2019; Stubberfield et al., 2019). Our data further confirms the same correlation and usefulness of WGS in a collection of bovine isolates. However, extended analysis with a larger number of test isolates would be desirable. Furthermore, all the above studies differed with respect to WGS databases used, AST clinical guidelines (CLSI/EUCAST), sample size and types and numbers of antibiotics examined. For instance, while our study examined 10 antibiotics from 8 antibiotic classes, another recent veterinary study on *E. coli* isolates from pig samples by Stubberfield et al. (2019) tested 9 antibiotics from 6 classes, with a larger sample size of 515. Even though the study by Stubberfield et al. (2019) may be more accurate due to the larger sample size, our study provides a novel insight into AMR correlation in bovine isolates, which often differ from isolates found in pigs with regards to strain type. A study by Feldgarden et al. (2019) examined 6,242 isolates including 294 *Campylobacter coli*, 476 *Campylobacter jejuni*, 47 *Escherichia coli*, and 5,425 *Salmonella enterica* isolates from human, animal and food sources, which is the largest sample size among the studies reported. However, the number of *E. coli* isolates was relatively low, and the animal sources were not specified. The only other study on bovine isolates was performed by Tyson et al. (2015), with only 76 *E. coli* isolates tested. Our study therefore provides an important insight on the agreement between genotypic and phenotypic AMR in bovine *E. coli*.

Several future approaches to AMR correlation analysis could be suggested. To improve correlation, isolates with three or more false positive/false negative results across three or more antimicrobial classes could be removed from the isolate group to exclude isolates where potential testing errors, sample discrepancies or other confounding factors may be at play. This approach was utilised previously to improve correlation in a similar study by Feldgarden et al. (2019). Isolates which are resistant to certain antimicrobial compounds but lack related AMR genes, i.e., false negative isolates, could be further examined and the reason for their resistance, such as encoding a novel AMR gene or an efflux pump, could be determined. Alternative methods of AST such as MIC by VITEK may yield AST results which would result in a better correlation with AST-WGS due to the reduction of unavoidable human error associated with manual methods.

To implement AST-WGS as a routine diagnostic test, standardisation of the DNA extraction, library preparation and bioinformatics analysis process would be necessary. There would be a need for harmonisation between laboratories and internationally, a huge task given the diversity in methods of bioinformatics analysis currently (Moran-Gilad, 2017). Although WGS may not be a faster method, the main advantage of this method over phenotypic methods of AST is that WGS provides much more information on each isolate tested. Not only can the exact gene(s) that confer resistance in isolates be identified, other useful information can be extracted from the sequence, such as the potential virulence of the isolates. Their relatedness to other isolates that were previously sequenced can be useful in public health situations where outbreaks occur and for outbreak investigations and epidemiological tracebacks. In hospital settings, WGS could be used to differentiate between true infection and environmental contamination, which is not currently possible using phenotypic methods (Chapman et al., 2020).

## Conclusion

The aim of this project was to determine if WGS could be used as an accurate predictor of AMR in bovine *E. coli* isolates. Results show promising agreement between AST-WGS and phenotypic AST. Resistance to tetracycline, florfenicol, gentamicin, trimethoprim-sulfamethoxazole and amoxicillin can be predicted using a genomic approach. However, more research is needed to study specific genes conferring resistance to amoxicillin-clavulanic acid, cephalothin, ciprofloxacin and marbofloxacin to improve correlation. Prediction of AMR phenotype requires careful curation of gene alleles conferring resistance, based on available literature.

Poor statistical correlation values may arise due to several factors such as 1) low numbers of resistant isolates in the sample set; 2) bovine *E. coli* having different genetic mechanisms of resistance when compared to human isolates; and 3) the large number of intermediately resistant isolates.

